# Single molecule fingerprinting reveals different amplification properties of α-synuclein oligomers and preformed fibrils in seeding assay

**DOI:** 10.1101/2021.08.09.455607

**Authors:** Derrick Lau, Chloé Magnan, Kathryn Hill, Antony Cooper, Yann Gambin, Emma Sierecki

## Abstract

The quantification of α-synuclein (α-syn) aggregates has emerged as a promising biomarker for synucleinopathies. Assays that amplify and detect such aggregates have revealed the presence of seeding-competent species in biosamples of patients diagnosed with Parkinson’s disease. However, multiple species such as oligomers and amyloid fibrils, are formed during the aggregation of α-synuclein and these species are likely to co-exist in biological samples and thus it remains unclear which species(s) are contributing to the signal detected in seeding assays. To identify which species can be detected in seeding assays, recombinant oligomers and preformed fibrils were produced and purified to characterise their individual biochemical and seeding potential. Here, we used single molecule spectroscopy to track the formation and purification of oligomers and fibrils at the single particle level and compare their respective seeding potential in an amplification assay. Single molecule detection validates that size-exclusion chromatography efficiently separates oligomers from fibrils. Oligomers were found to be seeding-competent but our results reveal that their seeding behaviour is very different compared to preformed fibrils in our amplification assay. Overall, our data suggest that even a low number of preformed fibrils present in biosamples are likely to dominate the response in seeding assays.

## Introduction

Parkinson’s Disease (PD) is characterised by the loss of neurons within the *substantia nigra pars compacta*, resulting in motor symptoms of tremors and rigidity to limbs, bradykinesia and stooped postures.[17] Although the etiology of PD is still not well understood, it is established that α-synuclein (α-syn) plays a central role in both idiopathic and familial forms of the disease. α-syn is a 14 kDa synaptic protein found across many cells of the central nervous system and is highly enriched in dopaminergic neurons.[1] Missense mutations of the α-syn encoding gene SNCA lead to an increased propensity of α-syn to aggregate into fibrils and are linked to familial forms of the disease.[22, 32, 47] Allele duplication of the SNCA gene also leads to PD.[19] It is hypothesised that α-syn aggregates are the catalysts that lead to the eventual formation of dense and compact intraneuronal inclusions known as Lewy bodies, the pathological hallmark of PD.[44] Dementia with Lewy bodies (DLB) and multiple system atrophy (MSA) are also clinically related to PD with the presence of intra-cellular α-syn aggregates. The consistent presence of α-syn aggregates in these pathologies and the demonstrated ability of these aggregates to spread to other cells with prion-like behaviour, make pathogenic forms of α-syn both a promising biomarker of synucleinopathies and a likely pathogenic mechanism of disease progression. Recent developments, such as real time quaking induced conversion assay (RT-QuIC) or Protein misfolding cyclic amplification assay (PMCA) utilise the self-templated amplification of α-syn to detect the presence of α-syn aggregates in biofluids, in particular, cerebral spinal fluid (CSF). Using these assays, different groups have successfully identified patients with diverse synucleinopathies.[36, 40] Importantly, recent studies suggest that RT-QuIC could identify PD patients in the prodromal phase,[16] offering new hope to the community.

The further development of amplification methods and quantitative assays requires the use of well-defined standards that recapitulate the behaviour of the biological samples. Such standards have been difficult to identify when working with α-syn. For example, α-syn aggregates isolated from brains encompass a range of post translational modifications, including truncations,[2, 43] phosphorylation,[12] acetylation,[2] nitration[13] and ubiquitination.[50] These modified α-syn aggregates are too heterogeneous to become reliable standards. Preformed aggregates generated from recombinant proteins have been utilised, but even this material can be difficult to standardise between laboratories.[21, 30, 37, 47] One object of debate in using preformed aggregates has been the relevance of using α-syn fibrils compared to oligomers. Indeed, during the self-association process, both *in vitro* and *in vivo*, α-syn is thought to change from a mainly unstructured monomer to form small, soluble oligomers (consisting of 30-50 monomers). A structural switch to antiparallel β-sheet is required for the oligomers[49] to form protofibrils that can “grow” by recruiting and incorporating more α-syn monomers, generating long amyloid fibrils. The pathological relevance of α-syn oligomers and fibrils remains undecided. α-syn oligomers appear to be the more toxic species, at least in the short term, as they have been shown to bind and disrupt cellular membrane and participate in mitochondrial and lysosomal dysfunction.[9, 23, 26, 47, 48] Oligomers also affect proteasome activity,[10] vesicular trafficking or induce endoplasmic reticulum stress by toxic gain-of-functions.[14] On the other hand, α-syn fibrils constitute the majority of Lewy Bodies and participate in the sequestration of important cellular factors; such as mitochondria,[25] driving cell dysfunction and/or cell death. Early products from the aggregation process are partially reversible and α-syn fibrils could serve as a reservoir of oligomers.[7, 42] Because of the difference in biological activity, the question of whether oligomers or fibrils are the most relevant species to serve as biomarkers arose. The high sensitivity of the amplification (seeding) assays, in the low femtomolar range, makes this question more pressing, as even a minority species present in minute quantities could dominate the amplification outcome.

Here we use a newly developed single molecule fluorescence method to determine the “fingerprints” of α-syn oligomers and fibrils and assess their seeding potential in an amplification assay. Single molecule imaging is emerging as a powerful tool to study molecular dynamics with the ability to resolve microscopic events that are otherwise hidden from ensemble average measurements (see review[33]). In this study, we profiled α-syn aggregates as they diffuse freely in solution using a 3D printed confocal microscope. This plug-and-play device is designed to detect single aggregates labelled with the established fluorescent dye, thioflavin T (ThT). When ThT reactive species cross the small excitation/detection confocal volume (~1 fL), individual peaks of fluorescence are detected above background to produce fluorescence traces that can be analysed. In addition to counting aggregates, this method also determines the ThT reactivity (prominence) and apparent size (residence time) for each event. We used this fingerprinting approach to track the formation and purification of oligomers that are currently evaluated as standards for amplification assays.[21, 30] Our single molecule measurements revealed the rapid formation of oligomers and fibrils within 5 h of a standard aggregation reaction and demonstrated that size exclusion chromatography (SEC) was necessary to completely separate monomeric, oligomeric and fibrillar species of α-syn. We then compared purified α-syn oligomers to sonicated α-syn fibrils, before and after an isothermal incubation to assess their amplification potentials. Our single molecule data show very different amplification profiles and seeding propensity for purified oligomers and fibrils. Our results suggest that in heterogeneous samples, a positive response in RT-QuIC assays would be mainly driven by the presence of fibrils.

## Materials and Methods

### Expression and purification of wild-type human α-syn

The plasmid pT7-7 (Addgene 36046)[29] coding for the human α-synuclein wild type (α-syn) was transformed into *E. coli* BL21 Rosetta (DE3, pLysS RARE) for expression in Luria-Bertani medium containing ampicillin (100 μg/mL) and chloramphenicol (34 μg/mL) at 37°C. Protein expression was induced with IPTG (1 mM) at an optical density (600 nm) of 0.5 and allowed to proceed at 18°C for 16 hours with shaking. Cells were harvested with centrifugation, resuspended in cold lysis buffer (25 mM Tris, pH 8, 0.02% w/v NaN_3_, Complete protease inhibitor, Roche, 04693132001) and lysed using the CF cell disruptor (Constant Systems Ltd) at 20 kPSI. Lysate was supplemented with EDTA (10 mM) and incubated at 90°C for 20 min to precipitate bacterial proteins. Lysate was clarified by centrifugation (Thermo Fisher Scientific, SS-34 rotor, 19000 rpm, 30 min, 4°C). The supernatant was retained and supplemented with streptomycin sulfate (10 mg/mL, Sigma Aldrich, S6501-25G), stirred for 20 min at 4°C and then centrifuged at 19000 rpm (SS-34 rotor, 20 min, 4°C) to recover the clarified supernatant. This incubation and centrifugation steps were repeated with 20 mg/mL and 30 mg/mL of streptomycin sulfate. The clarified supernatant was then incubated with ammonium sulfate (0.4 mg/mL) for 30 min at 4°C and centrifuged at 13600 rpm, 20 min, 4°C (SS-34). The pellet was resolubilised in buffer A (20 mM Tris, pH 7.7, 0.02% w/v NaN_3_) and dialysed overnight at 4°C (Thermo Fisher Scientific, 68700). Dialysed solution was filtered (0.22 μm) and further purified by anion exchange chromatography using 2 HiTrap Capto Q ImpRes columns (Cytiva, 17547055). The column was equilibrated with buffer A before injecting the sample at 1 mL/min. α-syn eluted at approximately 175 mM NaCl using a linear 300 mL gradient from 0 to 1 M NaCl in buffer A (3 mL/min). Fractions containing α-syn were identified using reducing SDS-PAGE, combined, and concentrated using Amicon Ultra 15 filters (Merck, UFC901024) for size exclusion chromatography. Elution was performed using a HiLoad 16/600 Superdex 200 column (GE Healthcare, 28989335) equilibrated with buffer A at 1 mL/min. Purity was assessed using reducing SDS-PAGE and samples were concentrated to > 343 μM (4.9 mg/mL). Protein concentration was determined spectroscopically at 280 nm absorbance with an extinction coefficient of 5960 M^−1^cm^−1^. Purified α-syn was aliquoted, flash frozen with liquid nitrogen and stored at −80°C. The yield was 4.2 mg/g of cell mass.

### Preparation of α-syn oligomers

Generation of α-syn oligomers was based on a protocol by Kumar et al.[21, 30] and Rösener et al.[35] Purified α-syn were lyophilised with the LyoQuest freeze dryer (Telstar) or evaporated at RT with the SpeedVac SC250 (Thermo Fisher Scientific) under vacuum overnight. Lyophilised proteins were solubilised in phosphate buffer saline (PBS, 2 mM KH2PO4, 10 mM Na2HPO4, 2.7 mM KCl and 137 mM NaCl, pH 7.4) to a final α-syn concentration of 12 mg/mL (830 μM) and incubated at 37°C for 5 h in a vortexer, shaken horizontally at 900 rpm in a 2 mL cryovial. An aliquot of the reaction mixture was withdrawn at different timepoints for single molecule measurements, flash frozen in liquid nitrogen and stored at −80°C until measurement. Approximately 500 μL of aggregation reaction at 5 h were frozen for SEC to purify α-syn oligomers. Five individual α-syn assemblies were performed across two different days. Products of the 5 h reactions were thawed at room temperature and centrifuged at 18000 g, 10 min, 4°C (Beckman Coulter F301.5 rotor) to remove large aggregates. Pellets were resuspended in PBS and kept for negative staining transmission electron microscopy (EM). The α-syn oligomers and free α-syn from the supernatant were separated using a Superdex 200 Increase 10/300 GL (GE Healthcare, 28990944) equilibrated with PBS. The concentration of α-syn in the different fractions were determined using gel densitometry (see Material and methods) and were estimated to contain 0.2 μM to 3.1 μM α-syn (monomer equivalent). Afterward, ThT (10 μM) and monomeric α-syn (30 μM) were added to the different fractions to stain for ThT^+^ (ThT reactive) species, prevent dissociation and binding to the surfaces, respectively. Negative controls of monomeric α-syn returned with an average of 1.4 events per trace despite the filtering of the α-syn stock through a 100k MWCO prior to use (Figure 4A). We believe this low-level detection comes from non-specific interaction of ThT with reagents trapped in the gel column as similar values were reflected in fractions in the void volume (elution volume 0-7 mL, Figure 4A)

### Single molecule analysis of aggregation kinetics

Aliquots withdrawn at different timepoints of the oligomer preparation were diluted by 20-fold in PBS containing 10 μM of ThT (Sigma Aldrich, T3516-5G), loaded to a custom polydimethylsiloxane (PDMS) plate adhered to a glass coverslip (ProSciTech, G425-4860) and observed using the inverted 3D printed confocal microscope, “AttoBright”, equipped with a 450 nm laser and water immersion 40x/1.2 NA objective (Zeiss).[6] Emitted fluorescence from ThT was filtered by a dichroic mirror (488 nm) and a long-pass filter (500 nm) before focusing onto a single photon avalanche diode (Micro Photon Devices). Fluorescence spectroscopy traces were recorded for 100 sec/trace in 10 ms bins and analysed using a custom python script (see Supplementary Information) on Spyder version 4.0.1 to extract information of large intensity bursts (peaks) in the fluorescent traces in an automated and unbiased manner.

The analysis script reports on:

1. The prominence of each burst. This represents the intensity of a peak corrected for background fluorescence. The prominence of each peak was determined by extending a horizontal line from the peak maxima to the left and right to intersect the raw signal. The lowest intensity value between the peak maxima and the new intercepts forms the base of a peak. Prominence is then defined as the difference of the lowest of the two intensity values of the bases and the peak maxima intensity.
2. The residence time or the full width half maximum (FWHM) of a burst. Equivalently, it is the difference of two time points in which the intensity values are at 50% of the prominence.
3. The total intensity of each fluorescence burst, as the sum of intensities with the upper and lower limit defined by the FWHM to estimate the area under the curve (AUC).

We determined that the FWHM of a peak provided a more reliable metric compared to using 90% of prominence or 100% (full width). This is because the latter gave rise to artificially large residence times, compounded by the fact that a Gaussian peak stretches to infinity in which peak bases can never be defined. Thus, the total intensity currently reported is an underestimation of the true intensity of each burst.

### Amplification of α-syn oligomers

SEC eluted fractions (in PBS) were mixed with filtered monomeric α-syn WT (30 μM) and ThT (10 μM) and were incubated at 55°C for 5 h in a PCR machine (Bio-rad, C1000) for amplification.[3] Fractions comprised of ≥ 76% of its original volume were mixed with α-syn and ThT to reach a final volume of 20 μL per reaction. Samples were loaded without dilution onto the PDMS plate to record fluorescence on AttoBright. Fluorescence was recorded in 100 sec/trace, 10 ms bins and was analysed using the custom script as previously described. Fractions that revealed significant positive increase in the number of ThT peaks after amplification were designated as fractions containing α-syn oligomers. These fractions were reproducibly identified across different SEC experiments and traces were pooled together for analysis.

### Production and sonication of α-syn preformed fibrils

Human α-syn WT preformed fibrils (PFFs) were generated by incubating a solution of monomeric α-syn WT (208 μM) in PBS at 45°C with shaking (500 rpm) for 72 h in the presence of a mini-stirrer. The solution was sonicated for 15 min at 12 h then every 24 h at room temperature in a water bath (Ultrasonics, FXP 14M) to induce fragmentation of the fibrils. PFFs were snap frozen in liquid nitrogen and stored at −80°C for future use. PFFs were thawed, loaded to a Microtube-50 AFA screw capped capsule (Covaris) and sonicated using a Covaris focused ultrasonicator (M220) for 10 min, 5 sec on/off at 75 Watts, 12°C. Samples were kept in the capsule at RT.

### Time course amplification of α-syn preformed fibrils and oligomers

Sonicated PFFs (2.6 nM) and three different elution fractions containing amplifiable oligomers (0.15 - 2.4 μM of α-syn) were mixed with α-syn WT filtered through a 100 k MWCO membrane (30 μM) and ThT (10 μM) in PBS. Reactions were aliquoted for immediate fluorescence measurements on the Attobright and measurements after 2.5, 5, 7.5 and 24 h incubation at 55°C. Fractions seeded with oligomers were pooled during the analysis using the custom peak analysis script.

### Negative staining transmission electron microscopy

A copper grid (200 mesh, coated with carbon and formvar, Ted Pella, 01811) was cleaned by glow discharged and a small solution of samples; pellet and supernatant of the reaction mixture, oligomeric fractions from SEC, preformed and sonicated fibrils, were applied onto the grid and wicked dry. The grid was washed with a drop of MilliQ water and immediately wicked dry to minimise phosphate deposit. This was repeated two more times. The grid was then stained with a drop of uranyl acetate (2% w/v) and wicked dry immediately. This process was repeated twice, and the grid was air dried. Micrographs were collected using a FEI Tecnai G2 20 electron microscopes at 38000-fold magnification. Particle diameters were measured using ImageJ.

### Estimation of protein concentration

The concentration α-syn from the purification was estimated spectrophotometrically with extinction coefficient of 5930 M^−1^cm^−1^ at 280 nm. The protein concentration in SEC eluted fractions was spectroscopically too low for accurate quantification. Therefore, the concentration of α-syn was estimated using gel densitometry with known amount of α-syn WT (100-3200 ng, determined spectroscopically) loaded on reducing SDS-PAGE as standard. Protein bands were revealed with 1 h staining with SimplyBlue Safe stain (Thermo Fisher Scientific, LC6065) and overnight de-staining in water. The gel was imaged using the Chemi-doc system (Biorad) with Cy5.5 filter that provided a linear readout for interpolation. The concentration of α-syn in the eluted fractions containing oligomers were low, ranging 0.2 – 3.1 μM.

## Results

### Principle of single molecule profiling of α-syn aggregates

We and others previously have demonstrated that single molecule fluorescence assays were compatible with the use of thioflavin T (ThT) to observe and characterise human α-syn oligomers and larger soluble aggregates (Figure 1A).[3, 28, 40]. We have shown that single molecule detection enables a > 100,000-fold increase in sensitivity over traditional plate reader measurements for ThT. Traditionally, single molecule measurements are performed on expensive commercial microscopes or on home-made setups that are difficult to replicate between laboratories. To improve the access to this method, we designed a small device (”AttoBright”) to perform counting and characterisation of single aggregates; the device can be easily replicated by 3D plastic printing.[6] ThT-stained (ThT^+^) particles are detected on this “AttoBright” microscope as they diffuse across the confocal volume, producing large bursts of fluorescent intensity (event) in the trace (Figure 1A). α-syn aggregates containing β-sheets bind strongly to ThT while monomeric α-syn does not bind to ThT. Fluorescence traces can then be quantitatively analysed using an automated algorithm to report the number of ThT events per trace, as well as the prominence (burst intensity), residence time and total intensity (i.e. the sum of all intensities to calculate the area under the curve) of each event (Figure 1B). The prominence and total intensity report on the number of ThT reactive particles in the sample while the residence time, extracted using the full-width-half-maximum (FWHM, time difference taken at half prominence), shows changes in size or compactness of ThT labelled α-syn aggregates. Overall, the algorithm designed here can extract biophysical properties of individual α-syn aggregates to generate a profile or fingerprint of α-syn sub-species in the population.

**Figure 1.**
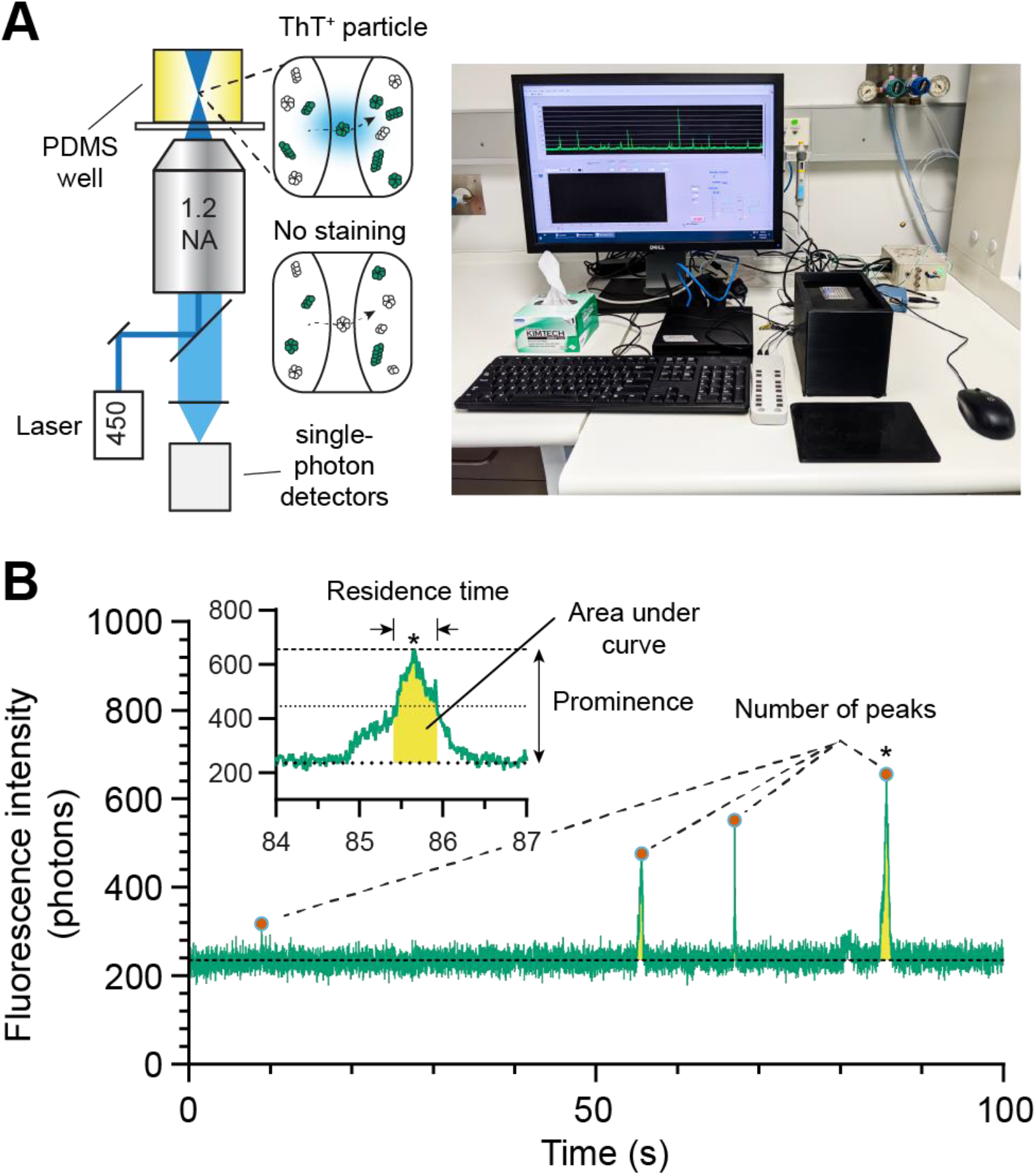
Single molecule fingerprinting for the characterisation of α-synuclein species. **A.** Left: schematic of the microscope setup. The inset shows ThT stained/dark α-syn oligomers or fibrils (green/white coloured particles) diffusing across the confocal volume. Monomeric α-syn and some assemblies do not bind ThT. Right: photograph of the microscope for recording the fluorescence traces. **B.**Characterisation of fluorescence traces. A fluorescence trace is analysed to report the total number of peaks (events) and individual peak’s prominence, residence time (full width half maximum) and area under the curve (yellow). The inset shows a region denoted by (*) in the trace.

### Rapid kinetics and heterogeneity in the early aggregation process

The AttoBright’s ability to detect and characterise α-syn aggregates at single molecule level provided the opportunity to examine the early molecular events of aggregation. Monomeric α-syn was incubated with shaking to produce oligomers.[21] Aliquots were taken at different time points during the reaction and diluted for single molecule fingerprinting experiments. Individual ThT^+^ peaks were detected as early as 2.5 h into the experiment (Supporting Figure 1). The number of events subsequently increased exponentially over time, generating a heterogeneous population of ThT^+^ particles (Figure 2B) up to a maximum of 51 events per 100 s trace after 5 h (Figure 2A-B). The short lag phase observed here is reasonable given the high starting concentration of α-syn (12 mg/mL) that would increase the probability of primary nucleation.

**Figure 2.**
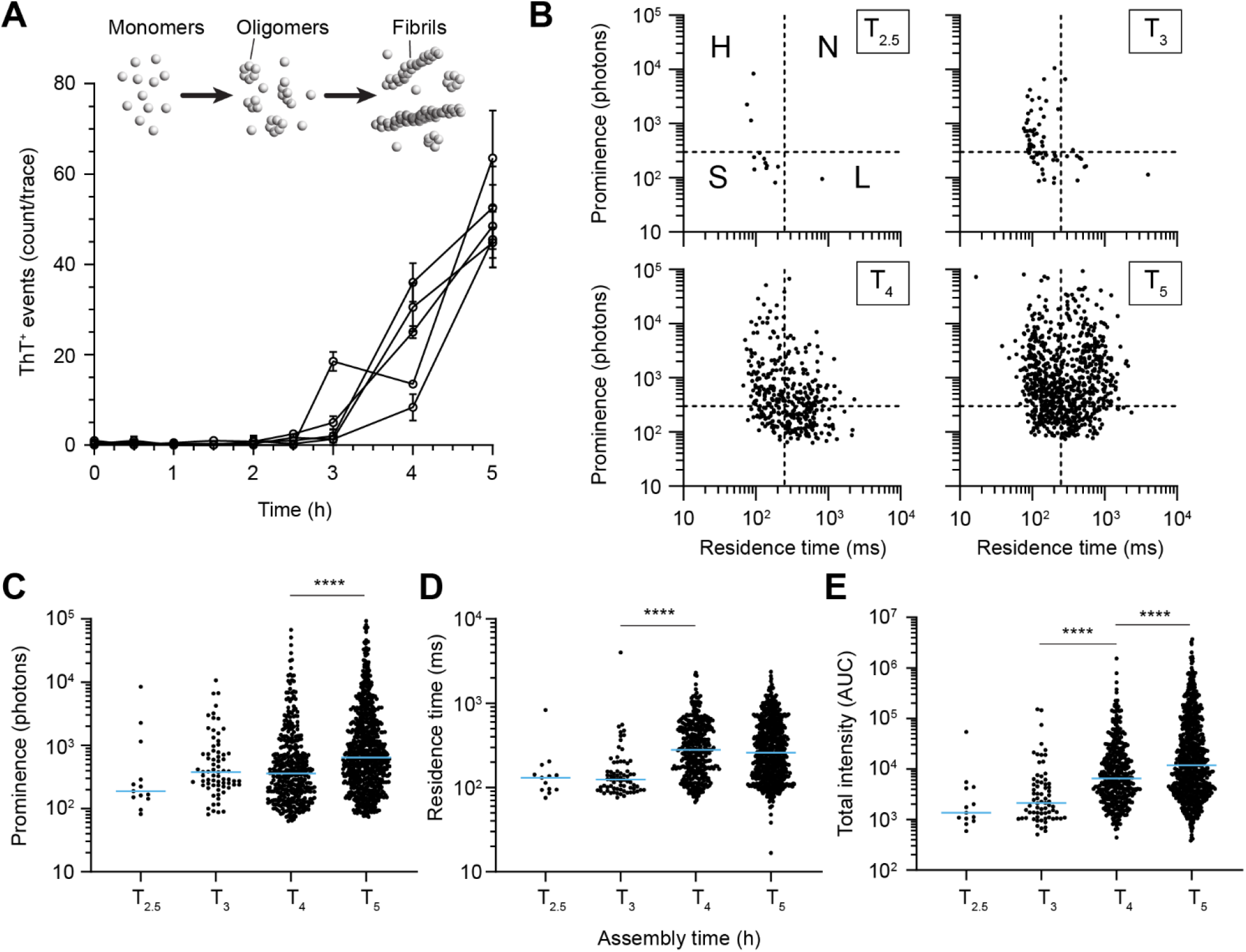
Single molecule aggregation kinetics and fingerprinting of species. **A.** Number of ThT^+^ species detected during the time course of α-syn aggregation. Each curve corresponds to an independent aggregation experiment. Error bars represent the standard deviation. **B.** Single molecule fingerprinting of ThT^+^ peaks detected. Peak prominence is plotted against its residence time from 2.5 h into the reaction. Four types of particles can be defined: small (S), high (H), long (L) and neutral (N). The cut-offs were set at 250 ms and 300 photons, on the x and y‐axis, respectively. **C-E.** Scatter plot comparing prominence (C), residence time (D) and total intensity (E). Blue line represents the median values. Each symbol represents an individual peak detected in the fluorescence traces generated from five independent experiments in panel A. Statistics used Kruskal-Wallis one-way ANOVA, p ≤ 0.0001 (****), non-significant (n.s.).

The prominence of each event was plotted against its respective residence time to examine population heterogeneity and provide a single-molecule profile (fingerprint) of the sample, as previously described.[3] Briefly, ThT^+^ particles were classed based on their distribution in each quadrant (Figure 2B) where events were defined as small (S, low prominence, short residence time), high (H, high prominence, short residence time), long (L, low prominence, long residence time) or neutral (N, high prominence, long residence time) with thresholds set at 300 photons and 250 ms respectively. Our data showed that S type particles were the first to be synthesised during the *in vitro* oligomer-inducing protocol. A total of 13 particles, pooled from 5 experiments, were recorded at 2.5 h. These species likely represent small α-syn oligomeric aggregates and are characterised by a median prominence of 188 photons, and median residence time of 130.4 ms (Figure 2C). The scatter profile showed significantly more particles after a further 30 min incubation and revealed the emergence of H type particles, from the possible conversion of S type oligomers. Interestingly, a small increase in prominence was noted with no change in the residence time (Figure 2B-C), suggesting that addition of monomers to protofibrils/oligomers assembly did not significantly increase their hydrodynamic radius. Beyond 3 h incubation, we observed the appearance of higher order assemblies of N and L type (Supporting Figure 1 and Figure 2) corresponding to larger particles with variable bound ThT ratios.

As expected, the total intensity increased over time (Figure 2E). Interestingly, our analysis shows a two-step assembly process. At early time points (< 3 h), increase in signal intensity is due to the increase in the number of events detected. Between 3-4 h, total intensity enhancement was contributed by a significant increase in residence time (median of 124 to 281 ms, Figure 2D) while prominence remained unchanged (median 378 to 360 photons, Figure 2C). In contrast, subsequent significant change in prominence became the main contributor to total intensity increase, at 4-5 h. At early time points, there appears to be a molecular event that makes the α-syn aggregate more ThT reactive. This could be a conformational folding event as β-sheets are required to recruit ThT.[28, 49] The L and N particles likely correspond to fibrils and fibrillar aggregates respectively and this was eventually verified by transmission electron microscopy (Figure 3B). We were unable to assert whether L and N particles were produced from the direct conversion of S or H particles, or both.

**Figure 3.**
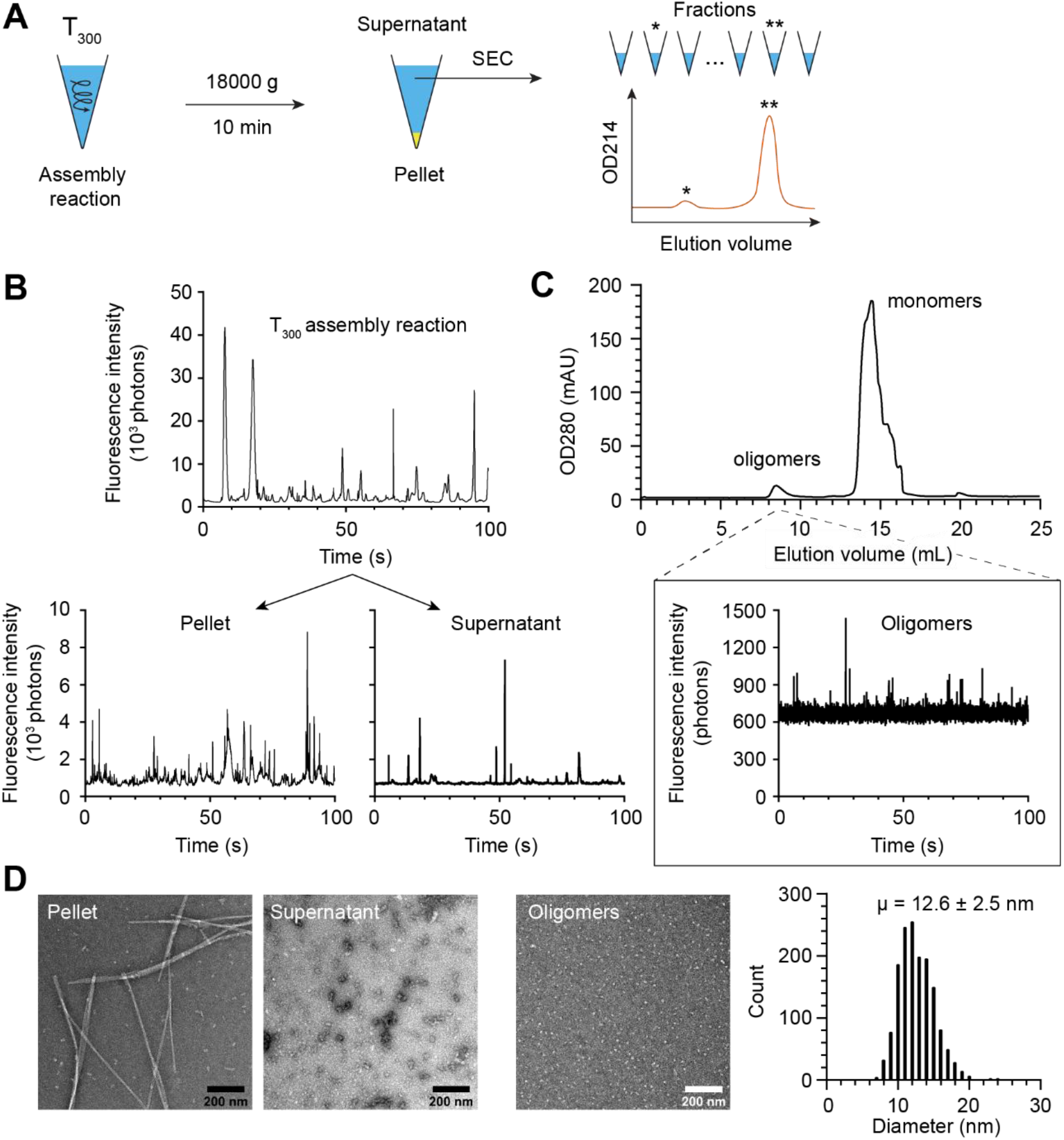
Isolation of α-syn oligomers. **A.** Schematic of the isolation protocol to isolate and detect ThT^+^ α-syn oligomers. After 5 h of aggregation, the reaction sample was centrifuged. The supernatant was further purified using size exclusion chromatography. Collected fractions were supplemented with monomers for amplification reactions and single molecule detection with ThT. **B.** Representative fluorescence traces of supernatant and pellet fraction with corresponding electron micrographs of each fraction shown in panel D. **C.** Gel filtration chromatogram identifying the elution peaks containing α-syn oligomers and monomers. A representative fluorescence trace corresponding to a fraction containing oligomers is shown. **D.** Electron micrographs of the pellet, supernatant and a SEC fraction containing oligomers. Histogram plot of the diameters of negative stained particles from the [9.2-9.7 mL] SEC fraction; 12.6 ± 2.5 nm; 1544/2 (mean ± standard deviation; number of particles/fractions stained). Scale bar is 200 nm.

Overall, these observations agreed with the α-syn nucleation cascade hypothesis that aggregation is a stepwise process driven by conversion of monomeric α-syn to small seeds (S-type) that will catalyse the formation more ThT reactive particles (H-type), possibly protofilaments, eventually forming fibrils and fibrillar aggregates (L or N-type).[27, 38, 49] Importantly, these data demonstrate that *in vitro* synthesis generates a heterogenous population of particles in the early aggregation process. Specifically, we show that a significant number of fibrils are produced within the first 5 h. These data emphasise the importance of performing additional steps to remove fibrils if a study makes explicit use of α-syn oligomers only.

Next, we were interested in isolating the small sized ThT species (S and H), to compare their seeding potential with fibrils. To do so, we first removed the large fibrils and aggregates and then removed the free monomers using a two-step purification protocol prescribed by Kumar et al[21] (Figure 3A).

### Removal of α-syn fibrils is incomplete by centrifugation

In brief, we centrifuged the 5 h reaction mixture at high speed (18,000 g, 10 min) to separate fibrils and smaller oligomers/aggregates. As expected, single molecule fingerprinting of the pellet showed slow diffusing ThT^+^ species (broad, high intensity events), in agreement with the presence of large fibrils observed under EM (Figure 3D). Single molecule analysis of the supernatant revealed faster diffusing particles (Figure 3B), and EM showed a heterogeneous population of small protofibrils and oligomers (Figure 3D). However, the high sensitivity of our single molecule assay revealed that slow diffusing species were still present in the supernatant, indicating that centrifugation was insufficient to fully remove mature fibrils and purify the smaller S and H oligomers (Figure 3B).

### Coupling single molecule detection on size exclusion chromatography demonstrates that isolation of α-syn oligomers is efficient and highly reproducible

Next, we coupled our single molecule detection system with size exclusion chromatography to visualise the isolation of oligomers and validate that all residual fibrils were removed. To do this, we simply collected all eluted fractions and characterised the number and size of aggregates by single molecule fingerprinting. The SEC chromatogram revealed two elution peaks: at 8.5 mL and 14.5 mL corresponding to oligomers and monomers in agreement to what has previously been reported (Figure 3C).[21, 31, 37]. As shown in Figure 4A, we identified a cluster of ThT^+^ events averaging 5.5 events per trace at the first elution peak in the SEC chromatogram, which corresponds to α-syn oligomers, according to previous publications.[21] In contrast, the low number of ThT^+^ species in the α-syn monomers peak (14-16 mL) was comparable to background level, confirming that α-syn monomers were, as expected, not ThT reactive. Surprisingly, we did not detect the slower diffusing fibrillar ThT species that were originally present in the supernatant in any eluted fractions (compare Figure 3B-C). We ascribed this discrepancy to the poor stability of α-syn protofibrils and oligomers[31, 42] with the dilution effect of gel filtration facilitating their disassembly.[7] Chromatograms obtained from four separate experiments showed identical profiles, demonstrating robust reproducibility.

**Figure 4.**
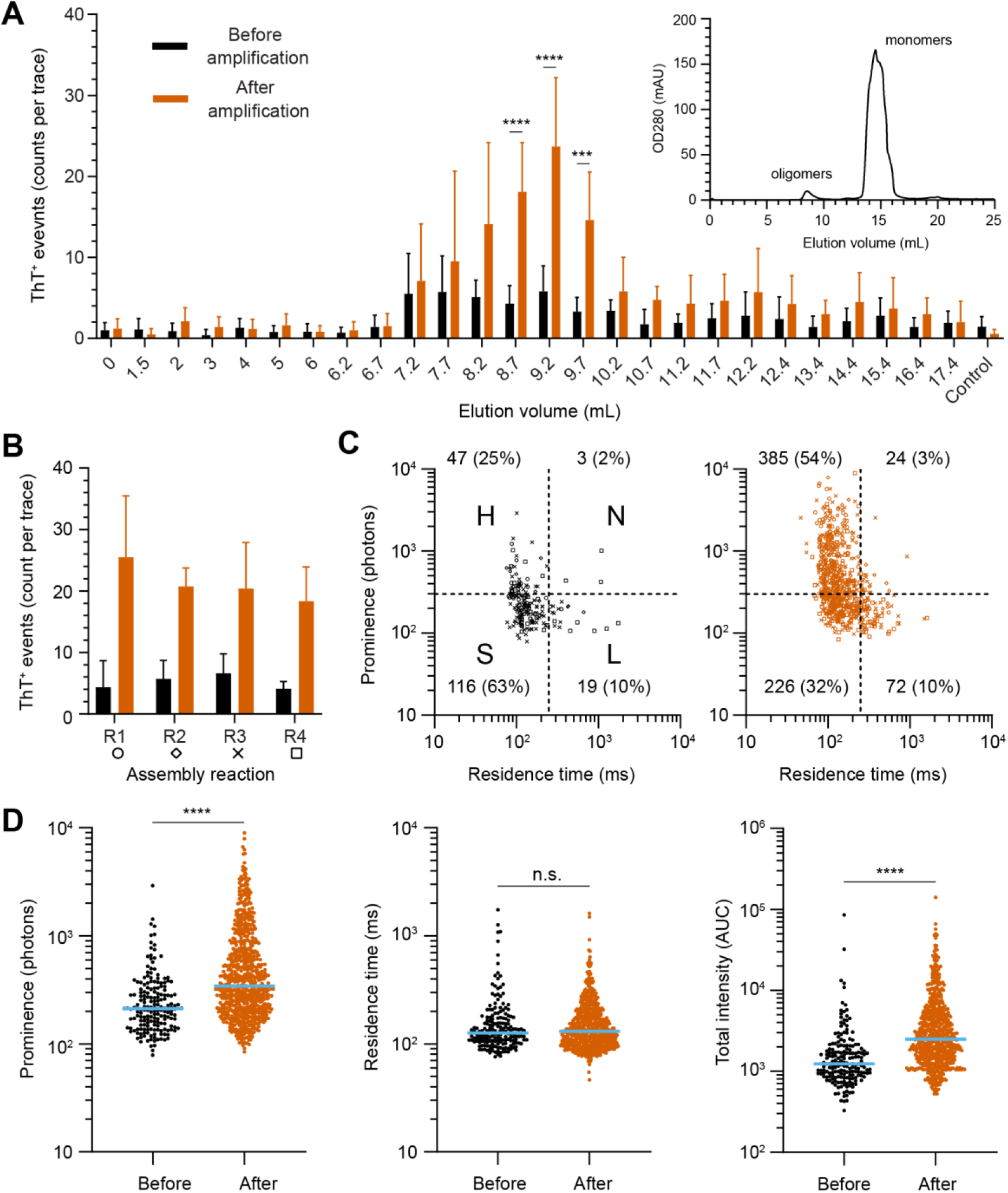
Single molecule characterisation of α-syn oligomers. **A.** Quantification of ThT^+^ events detected in each fraction after size exclusion, before (black) and after an amplification step (red) in PBS. Negative control is monomeric α-syn and ThT in PBS. A gel filtration profile is shown in the inset for reference. Error bars represent the standard deviation. This colour scheme is applied across all figures in this study. **B.** Bar graph of the number of ThT^+^ events detected in fractions containing oligomers (elution volume 7.7-10.2 mL) collected from four assembly reactions (R1-R4). **C.** Single molecule fingerprinting of oligomers, before (left) and after amplification (right), is obtained by plotting the prominence against residence time. Each symbol represents an individual event with the shape corresponding to R1-R4 coded in panel B. Quadrant numbers report the number of particles in each quadrant and abundance. **D.** Logarithmic scatter plot comparing peak prominence, residence time and area under the curve before and after amplification. Each symbol represents an individual event (N = 34/34 traces, before/after amplification). The median value is highlighted in blue. Statistics used Mann-Whitney t-test, p ≤ 0.0001 (****), non-significant (n.s.).

EM observation of the 8.5 mL fraction (centre of the oligomers peak) revealed particles with a narrow distribution of 12.4 ± 2.5 nm in diameter (Figure 3D). The ultrastructure and dimension of these particles is within range of previous measurements of 12 nm.[21] Our single molecule fingerprinting reveals that a fraction of the oligomers appears as ThT positive species and that their profiles correspond exclusively to S and H-type particles. In conclusion, gel filtration is a valid and very reproducible approach in isolating α-syn oligomers (categorised as S and H fingerprints) as identical elution profiles were observed across purification of different batches of α-syn aggregation experiment.

### Amplification potential of α-syn oligomers

We speculated that oligomers isolated in the early steps of the aggregation process would be seeding competent given they are hypothesised to be aggregation intermediates. To test this, we used an amplification assay where we added monomeric α-syn to all the SEC fractions and measured changes in ThT fluorescence after amplification. In this assay adapted for sensitive single molecule detection in small volumes, we use a single incubation step at 55°C in non-shaking conditions and measure the same sample 5 h later.[3] We observed that amplification greatly enhanced the number of ThT^+^ species in purified oligomers. As shown in Figure 4A, the increase in ThT^+^ reactivity was strictly limited to fractions containing α-syn oligomers (p < 0.002). In contrast, negative controls and monomeric α-syn displayed insignificant changes in ThT^+^ species. Amplification of the oligomeric fractions obtained from 4 independent chromatographic runs (R1-R4) showed that the number of events consistently increased by at least 3 times in the oligomeric fractions (Figure 4B). Before amplification, single molecule profiling of these pooled particles showed that S type particles (low prominence, low residence time) were the dominant species (63%, Figure 4C) followed by H type aggregates (25%). Importantly, large α-syn oligomers (> 250 ms residence time, 12%) were in low abundance.

The increase in the number of S type oligomers after the amplification step indicated the existence of dark (i.e., non-ThT reactive or has insufficient number of bound ThT molecules to overcome the background noise) seeding competent oligomers (Figure 4C). Concomitantly, we also observed an increase in the number of H particles (47 to 385) that represented the dominant species after amplification while the N and L populations remained unchanged. The isolated oligomers therefore amplified with a distinctive pattern where the increase in total intensity (peak area) was only contributed by an increase in ThT reactivity with limited change in its residence time (Figure 4D). The molecular signature of early oligomeric species can thus be best described by an upward transition of S to H particles (Figure 4C).

### The amplification profile of α-syn is distinct between sonicated fibrils and oligomers

Next, we sought to characterise the amplification profile α-syn PFFs and compare it to discern if they have a distinct *in vitro* signature from that of amplified α-syn oligomers. PFFs were created *in vitro* as described,[6] producing fibrils of 15.2 ± 2.1 nm in diameter (mean ± SD, Figure 5A) with longitudinal twists. This diameter is consistent with the “high salt strain” reported by Bousset et al.[5] To reduce the length of each fibril and make them amenable to spectroscopy measurement, PFFs were sonicated into smaller objects ranging from 10 to 90 nm (Figure 5A). Fluorescence measurement identified these particles as fast diffusing objects (< 250 ms) with variable ThT reactivity (Figure 5B). Overall, in the absence of supplementation of monomeric α-syn and heating (i.e. unamplified), the sonicated PFFs displayed a single molecule profile similar to the purified α-syn oligomers (compare Figure 4C with Figure 5C). In contrast, a 5 h amplification of sonicated PFFs produced a right-ward lateral shift (longer residence time) in the profile where L and N particles became predominant, accounting for 78% of the detected species. This molecular fingerprint contrasts strongly with the amplification signature of α-syn oligomers that was previously characterised as an upward shift in ThT reactivity with no change in residence time.

**Figure 5.**
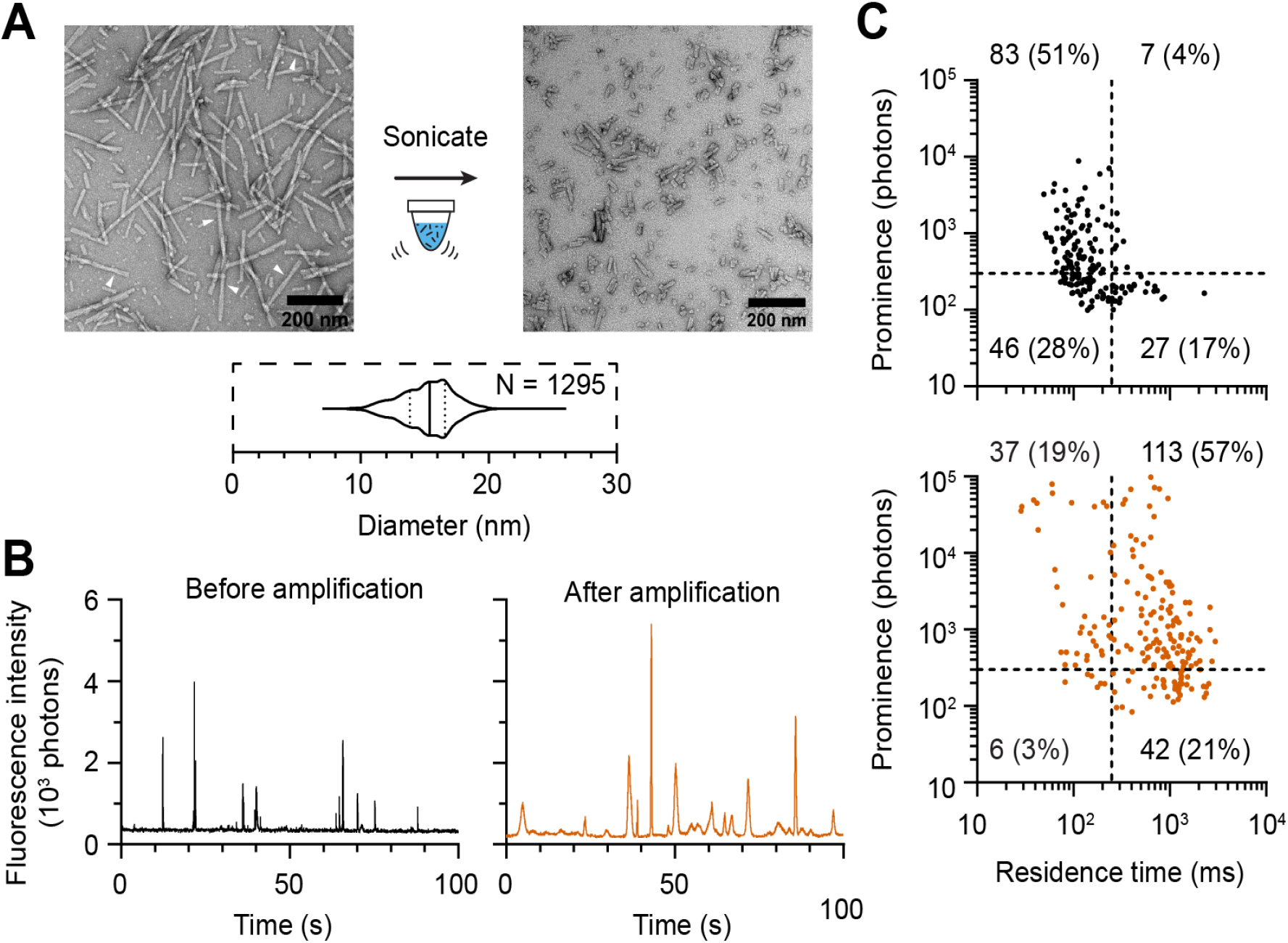
Single molecule fingerprinting of sonicated α-syn fibrils. **A.** Negative stained EM images of human α-syn fibrils before and after sonication. White arrows point at filamentous twists. Violin plot of tubular diameters of the fibrils. Scale bar is 200 nm. **B.** Representative fluorescence traces of sonicated fibrils before (black) and after a 5 h amplification (red). **C.** Single molecule fingerprinting of sonicated mature human α-syn fibrils before and after amplification (red). Each symbol represents an individual event in the fluorescence traces. Quadrant numbers represent the number of ThT events. Data from three independent experiments.

We expanded our analysis by performing amplification kinetics to examine the evolution of species over time in seeded experiments. In longitudinal analysis, we noted that, again, the increase in total fluorescence upon α-syn oligomers seeding occurs through an increase in ThT reactivity with little change in residence time over the course of 24 h (Figure 6A). This reemphasises that remodelling to enriched β-sheet protofibrils drives the process rather than elongation that would increase hydrodynamic radius. In contrast, the incubation of sonicated PFFs promoted rapid elongation of fibrils as evident from the seven-fold increase in median residence time after 2.5 h of incubation. Incubation of 2.5 h was sufficient to induce a 4-fold increase in the number of total ThT events when seeded with oligomers whereas no significant generation of new ThT species were observed when seeded with sonicated fibrils (Supporting Figure 4A). Again, this points to the likelihood that oligomers are first converting into ThT^+^ species through a refolding event to become structurally more β-sheets enriched, while fibrils amplify by elongating. Furthermore, seeding experiments with oligomers had an increased proportion of S and H particles within 2.5 h and maintained close to a 1 to 1 ratio (S:H) throughout the prolonged incubation (Supporting Figure 4B). In contrast, PFFs saw a redistribution of the population towards slower diffusing L and N particles.

**Figure 6.**
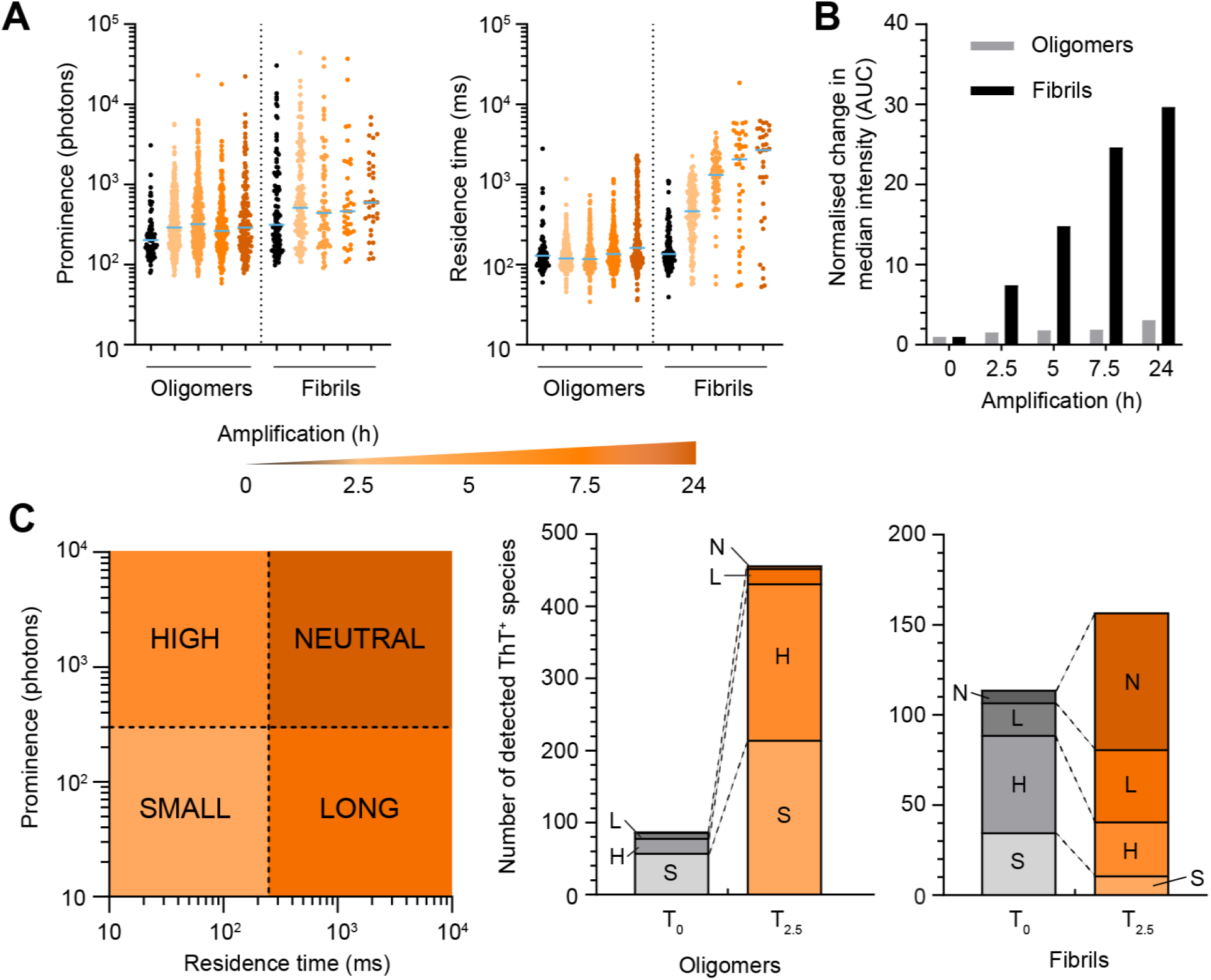
Time course amplification assay comparing α-syn oligomers and sonicated preformed fibrils. **A.** Time course measurements of oligomers or fibrils amplified for 2.5, 5, 7.5 and 24 h (orange to red) in the presence of excess α-syn monomers and ThT in PBS. Scatter dot plots comparing the prominence and residence time. Each symbol represents an individual ThT event. Data were collected from two independent experiments. The median value is highlighted in blue. **B.** Bar graphs comparing the fold-change in median AUC of fibrils and oligomers of Panel A. Value were normalised to the median total intensity collected at time 0. **C.** Time comparison on the number of ThT^+^ species of panel A detected from seeding experiment using α-syn oligomers or sonicated PFFs before and after 2.5 h incubation. Each type of ThT positive species is coded by different shades of red.

### Sonicated preformed fibrils amplify more efficiently than oligomers

The apparent increase in total ThT fluorescence in the amplification assay is much larger for preformed fibrils than for oligomers. When we calculate the increase in median ThT fluorescence of each particle over time, we find that the fluorescence signal has increased by 30-fold when the reaction was seeded by PFFs while seeding by purified oligomers only resulted in a 3-fold increase in fluorescence in a period of 24 h (Figure 6B). The enhanced efficiency of PFFs to amplify is even more dramatic when comparing the effective concentration of monomeric α-syn where seeding experiment with PFFs used at 2.6 nM (monomer equivalent) being three-orders of magnitude lower than oligomers (2 μM monomer equivalent). Oligomers can seed to form small protofibrils but are significantly slower in polymerisation compared to mature fibrils. We conclude that fibrils have significantly higher amplification (seeding) potential in comparison to its earlier oligomeric precursors. This observation is consistent with previous work that demonstrated α-syn fibrils folded β-sheet more rapidly than oligomeric mutants.[47]

## Discussion

One of the main challenges of performing *in vitro* studies to model for Parkinson’s disease is the heterogeneity of α-syn preparations which can have direct consequences on experimental reproducibility. Increasingly, observations that amyloid strains, including in α-syn fibrils, play decisive roles in disease presentation and progression, further increase this concern. The notion of “oligomers” itself remains ill-defined and has been used to describe sub-species of different sizes, shapes, and properties of α-syn aggregates depending on the preparation protocol used. For these reasons, a better definition of the sub-species and a standardisation of protocols is needed so that data can be more robustly reproduced and compared. Significant efforts in the recent years have been directed to standardise or characterise α-syn oligomers or PFFs. Approaches such as electron microscopy, atomic force microscopy, dynamic light scattering, and circular dichroism have been used for the physical characterisation of α-syn sub-species while biochemical assessments include proteinase digestion and the examination of aggregation kinetics.[9, 21, 22, 47] Most of these methods perform measurements in bulk and thus do not easily reveal the heterogeneity of the preparations. We have established that the detection limit for PFFs using AttoBright was 1 pM monomer equivalent.[3] This sensitivity enabled us to compare the ability of different steps to separate α-syn into different species (monomer, oligomer, fibril etc.). After centrifugation of a 5 h aggregation reaction (18,000 g for 10 min), many fibrils were still present in the supernatant and dominated the fluorescence time trace (Supporting Figure 2). Size exclusion chromatography was the preferred approach to obtain oligomers as it excluded both monomeric and fibrillar α-syn. Complete removal of fibrils (within the detection limit) was indeed observed by EM and confirmed via single molecule fingerprinting after size exclusion. We also explored the influence of lyophilisation on the stability of α-syn oligomers and their seeding ability. Reconstitution of lyophilised, purified α-syn oligomers resulted in a significant increase in the number of ThT^+^ particles, but in an apparent reduction in the amplification potential of the oligomers (Supporting Figure 5).

Here we provided single molecule profiles of the different α-syn sub-species and showed their relative distribution at different steps of their preparation and purification. We first tracked α-syn aggregation kinetics, and observed that self-assembly occurred rapidly, with ThT^+^ events detectable as early as 2.5 h into the incubation. Single molecule fingerprinting showed that small ThT^+^ species eventually transformed into slower diffusing protofibrils over time, producing a heterogeneous population of ThT^+^ amyloids. The resolution to track the evolution of ThT^+^ species over time lends support to the α-syn cascade hypothesis where α-syn monomers generate a population of oligomers that subsequently assemble into larger aggregates. The subsequent isolation of oligomers using gel filtration revealed that a fraction of these oligomers was indeed ThT positive. The estimated concentration of ThT reactive oligomers was ~0.6 nM by comparison with sonicated PFFs as our standard. This would imply that only < 0.03% of α-syn (of 2 μM) are incorporated into ThT^+^ oligomers. We subsequently identified another population of α-syn (~0.1% of 2 μM obtained from SEC) that became ThT reactive upon incubation with additional monomeric α-syn. We believe that these may correspond to dark oligomers that changed conformation to bind to ThT more efficiently. It could also be that these oligomers converted to short soluble protofibrils, thereby reaching the critical size for detection.[49]

Oligomers principally convert to form more fluorescent species (H-type), both during the initial aggregation process, as well as after purification. This increase in ThT fluorescence of individual aggregates likely stems from remodelling or elongation of the aggregate to form protofibril. Within a limited range, protofibril growth would occur without significantly affecting the residence time of the object but addition of subunits would increase the number of binding interfaces for ThT. Indeed, the single molecule fingerprint of sonicated fibrils was similar to α-syn oligomers even though the sonicated PFFs were more heterogeneous and often longer than the oligomers (12 ± 2 nm vs. 10-90 nm for the α-syn oligomers and sonicated fibrils, respectively). In the last phase of the kinetics, we observed a conversion of species from S/H to L/N types. This was similar to the amplification profile of the sonicated PFFs and differs from the amplification profile of the oligomers. Overall, these data support the recent observations by Ruggeri et al.[38] where the authors concluded that α-syn aggregation proceeds by an initial elongation of oligomers into single-strand protofibrils that then associate as “double-strand cross-section” protofilaments before forming mature fibrils.

The biological implication of toxicity between fibrils and oligomers remains controversial. On one hand, certain types of α-syn oligomers were demonstrated to trigger calcium influx induced cell death in SH-SY5Y cell lines.[7, 9] Similarly, the oligomers-forming mutants E35K and E57K, have higher affinities to interact with liposomes *in vitro* and a rat model of synucleinopathies have reported more cell deaths with these mutations.[47] On the other hand, fibrils are the accumulated species in Lewy bodies that are pathognomonic to PD with higher seeding potential for cell-cell transmission.[15, 24] These differing observations emphasise the need to correctly segregate and characterise each species prior to biological experiments. While these *in vitro* and *ex vivo* experiments have provided interesting insights in α-syn aggregation, the biological relevance of α-syn oligomers or fibrils in human biofluids remains an open question. We previously observed that α-syn aggregates amplified from CSF had a specific fingerprint compared to control [3] (albeit in different reaction conditions compared to this study). We hope that further definition of the α-syn species will inform on the biologically relevant assemblies that are found in biofluids.

## Conclusion

RT-QuIC and related methods are becoming tractable approaches for end-point diagnostic detection due to their high sensitivity to detect α-syn aggregates.[36, 39–41] However, these tools lack the ability to resolve features that may only be revealed at a single molecule level. The fluorescence approach described here is conceptually equivalent to RT-QuIC since it relies on ThT staining with an isothermal amplification step. However, our procedure does not involve mechanical shearing, compared to RT-QuIC [36]. In our hands, sonicated PFFs are far more seeding-competent than α-syn oligomers, at least at early time points of the amplification process. Therefore, it is very likely that the signal observed in RT-QuIC will be mainly driven by the presence of fibrils in a sample, at low concentration, even if α-syn oligomers were present. Furthermore, we demonstrated that isothermal amplification of sonicated PFFs produced very large species, resulting in a single molecule fingerprint that was distinct from amplified oligomers. Classifying α-syn sub-species based on their biophysical attributes adds an additional layer of granularity in the analysis whereby fingerprinting for different types of synucleinopathies may be obtained. This method of single molecule profiling can be applied to the detection and characterisation of other types of amyloids (e.g. Tau and prion proteins),[11, 20, 34, 46] study of other neurodegenerative diseases or the mechanistic investigation of putative cross-talks between α-syn/amyloid-β[18] and α-syn/Tau.[8, 18] ThT can fluorescently mark other large macromolecular assemblies such as fibrinogen[45] and DNA,[4] thus emphasising the importance of the isothermal incubation step with the appropriate monomeric substrate for specific detection of amyloids which can be easily optimised.

## Acknowledgements

This work was supported by grant MJFF-010267 from the Michael J Fox Foundation for Parkinson’s research and Shake it Up! Australia (to ES, YG and AC). AC received grant funding from the Australian Government. All authors have participated in writing the manuscript and have approved the submitted draft.

## References

1. Baba M, Nakajo S, Tu PH, Tomita T, Nakaya K, Lee VM, Trojanowski JQ, Iwatsubo T (1998) Aggregation of alpha-synuclein in Lewy bodies of sporadic Parkinson’s disease and dementia with Lewy bodies. Am J Pathol 152: 879–884

2. Bhattacharjee P, Öhrfelt A, Lashley T, Blennow K, Brinkmalm A, Zetterberg H (2019) Mass Spectrometric Analysis of Lewy Body-Enriched α-Synuclein in Parkinson’s Disease. J Proteome Res 18: 2109–2120 Doi 10.1021/acs.jproteome.8b00982

3. Bhumkar A, Magnan C, Lau D, Jun ESW, Dzamko N, Gambin Y, Sierecki E (2021) Single-Molecule Counting Coupled to Rapid Amplification Enables Detection of α-Synuclein Aggregates in Cerebrospinal Fluid of Parkinson’s Disease Patients. Angew Chem Int Ed Engl: Doi 10.1002/anie.202014898

4. Biancardi A, Biver T, Burgalassi A, Mattonai M, Secco F, Venturini M (2014) Mechanistic aspects of thioflavin-T self-aggregation and DNA binding: evidence for dimer attack on DNA grooves. Phys Chem Chem Phys 16: 20061–20072 Doi 10.1039/c4cp02838d

5. Bousset L, Pieri L, Ruiz-Arlandis G, Gath J, Jensen PH, Habenstein B, Madiona K, Olieric V, Böckmann A, Meier BHet al (2013) Structural and functional characterization of two alpha-synuclein strains. Nat Commun 4: 2575 Doi 10.1038/ncomms3575

6. Brown JWP, Bauer A, Polinkovsky ME, Bhumkar A, Hunter DJB, Gaus K, Sierecki E, Gambin Y (2019) Single-molecule detection on a portable 3D-printed microscope. Nat Commun 10: 5662 Doi 10.1038/s41467-019-13617-0

7. Cascella R, Chen SW, Bigi A, Camino JD, Xu CK, Dobson CM, Chiti F, Cremades N, Cecchi C (2021) The release of toxic oligomers from α-synuclein fibrils induces dysfunction in neuronal cells. Nat Commun 12: 1814 Doi 10.1038/s41467-021-21937-3

8. Castillo-Carranza DL, Guerrero-Muñoz MJ, Sengupta U, Gerson JE, Kayed R (2018) α-Synuclein Oligomers Induce a Unique Toxic Tau Strain. Biol Psychiatry 84: 499–508 Doi 10.1016/j.biopsych.2017.12.018

9. Danzer KM, Haasen D, Karow AR, Moussaud S, Habeck M, Giese A, Kretzschmar H, Hengerer B, Kostka M (2007) Different species of alpha-synuclein oligomers induce calcium influx and seeding. J Neurosci 27: 9220–9232 Doi 10.1523/jneurosci.2617-07.2007

10. Emmanouilidou E, Stefanis L, Vekrellis K (2010) Cell-produced alpha-synuclein oligomers are targeted to, and impair, the 26S proteasome. Neurobiol Aging 31: 953–968 Doi 10.1016/j.neurobiolaging.2008.07.008

11. Fiorini M, Iselle G, Perra D, Bongianni M, Capaldi S, Sacchetto L, Ferrari S, Mombello A, Vascellari S, Testi Set al (2020) High Diagnostic Accuracy of RT-QuIC Assay in a Prospective Study of Patients with Suspected sCJD. Int J Mol Sci 21: Doi 10.3390/ijms21030880

12. Fujiwara H, Hasegawa M, Dohmae N, Kawashima A, Masliah E, Goldberg MS, Shen J, Takio K, Iwatsubo T (2002) alpha-Synuclein is phosphorylated in synucleinopathy lesions. Nat Cell Biol 4: 160–164 Doi 10.1038/ncb748

13. Giasson BI, Duda JE, Murray IV, Chen Q, Souza JM, Hurtig HI, Ischiropoulos H, Trojanowski JQ, Lee VM (2000) Oxidative damage linked to neurodegeneration by selective alpha-synuclein nitration in synucleinopathy lesions. Science 290: 985–989 Doi 10.1126/science.290.5493.985

14. Gonçalves SA, Outeiro TF (2017) Traffic jams and the complex role of α-Synuclein aggregation in Parkinson disease. Small GTPases 8: 78–84 Doi 10.1080/21541248.2016.1199191

15. Hansen C, Angot E, Bergström AL, Steiner JA, Pieri L, Paul G, Outeiro TF, Melki R, Kallunki P, Fog Ket al (2011) α-Synuclein propagates from mouse brain to grafted dopaminergic neurons and seeds aggregation in cultured human cells. J Clin Invest 121: 715–725 Doi 10.1172/jci43366

16. Iranzo A, Fairfoul G, Ayudhaya ACN, Serradell M, Gelpi E, Vilaseca I, Sanchez-Valle R, Gaig C, Santamaria J, Tolosa Eet al (2021) Detection of α-synuclein in CSF by RT-QuIC in patients with isolated rapid-eye-movement sleep behaviour disorder: a longitudinal observational study. Lancet Neurol 20: 203–212 Doi 10.1016/s1474-4422(20)30449-x

17. Jankovic J (2008) Parkinson’s disease: clinical features and diagnosis. J Neurol Neurosurg Psychiatry 79: 368–376 Doi 10.1136/jnnp.2007.131045

18. Kayed R, Dettmer U, Lesné SE (2020) Soluble endogenous oligomeric α-synuclein species in neurodegenerative diseases: Expression, spreading, and cross-talk. J Parkinsons Dis 10: 791–818 Doi 10.3233/jpd-201965

19. Konno T, Ross OA, Puschmann A, Dickson DW, Wszolek ZK (2016) Autosomal dominant Parkinson’s disease caused by SNCA duplications. Parkinsonism Relat Disord 22 Suppl 1: S1–S6 Doi 10.1016/j.parkreldis.2015.09.007

20. Kraus A, Saijo E, Metrick MA, 2nd, Newell K, Sigurdson CJ, Zanusso G, Ghetti B, Caughey B (2019) Seeding selectivity and ultrasensitive detection of tau aggregate conformers of Alzheimer disease. Acta Neuropathol 137: 585–598 Doi 10.1007/s00401-018-1947-3

21. Kumar ST, Donzelli S, Chiki A, Syed MMK, Lashuel HA (2020) A simple, versatile and robust centrifugation-based filtration protocol for the isolation and quantification of α-synuclein monomers, oligomers and fibrils: Towards improving experimental reproducibility in α-synuclein research. J Neurochem 153: 103–119 Doi 10.1111/jnc.14955

22. Lashuel HA, Petre BM, Wall J, Simon M, Nowak RJ, Walz T, Lansbury PT, Jr. (2002) Alpha-synuclein, especially the Parkinson’s disease-associated mutants, forms pore-like annular and tubular protofibrils. J Mol Biol 322: 1089–1102 Doi 10.1016/s0022-2836(02)00735-0

23. Ludtmann MHR, Angelova PR, Horrocks MH, Choi ML, Rodrigues M, Baev AY, Berezhnov AV, Yao Z, Little D, Banushi Bet al (2018) α-synuclein oligomers interact with ATP synthase and open the permeability transition pore in Parkinson’s disease. Nat Commun 9: 2293 Doi 10.1038/s41467-018-04422-2

24. Luk KC, Song C, O’Brien P, Stieber A, Branch JR, Brunden KR, Trojanowski JQ, Lee VM (2009) Exogenous alpha-synuclein fibrils seed the formation of Lewy body-like intracellular inclusions in cultured cells. Proc Natl Acad Sci U S A 106: 20051–20056 Doi 10.1073/pnas.0908005106

25. Mahul-Mellier AL, Burtscher J, Maharjan N, Weerens L, Croisier M, Kuttler F, Leleu M, Knott GW, Lashuel HA (2020) The process of Lewy body formation, rather than simply α-synuclein fibrillization, is one of the major drivers of neurodegeneration. Proc Natl Acad Sci U S A 117: 4971–4982 Doi 10.1073/pnas.1913904117

26. Mazzulli JR, Xu YH, Sun Y, Knight AL, McLean PJ, Caldwell GA, Sidransky E, Grabowski GA, Krainc D (2011) Gaucher disease glucocerebrosidase and α-synuclein form a bidirectional pathogenic loop in synucleinopathies. Cell 146: 37–52 Doi 10.1016/j.cell.2011.06.001

27. Mehra S, Sahay S, Maji SK (2019) α-Synuclein misfolding and aggregation: Implications in Parkinson’s disease pathogenesis. Biochim Biophys Acta Proteins Proteom 1867: 890–908 Doi 10.1016/j.bbapap.2019.03.001

28. Naiki H, Higuchi K, Hosokawa M, Takeda T (1989) Fluorometric determination of amyloid fibrils in vitro using the fluorescent dye, thioflavin T1. Anal Biochem 177: 244–249 Doi 10.1016/0003-2697(89)90046-8

29. Paleologou KE, Schmid AW, Rospigliosi CC, Kim HY, Lamberto GR, Fredenburg RA, Lansbury PT, Jr., Fernandez CO, Eliezer D, Zweckstetter Met al (2008) Phosphorylation at Ser-129 but not the phosphomimics S129E/D inhibits the fibrillation of alpha-synuclein. J Biol Chem 283: 16895–16905 Doi 10.1074/jbc.M800747200

30. Paslawski W, Lorenzen N, Otzen DE (2016) Formation and Characterization of α-Synuclein Oligomers. Methods Mol Biol 1345: 133–150 Doi 10.1007/978-1-4939-2978-8_9

31. Pieri L, Madiona K, Melki R (2016) Structural and functional properties of prefibrillar α-synuclein oligomers. Sci Rep 6: 24526 Doi 10.1038/srep24526

32. Polymeropoulos MH, Lavedan C, Leroy E, Ide SE, Dehejia A, Dutra A, Pike B, Root H, Rubenstein J, Boyer Ret al (1997) Mutation in the alpha-synuclein gene identified in families with Parkinson’s disease. Science 276: 2045–2047 Doi 10.1126/science.276.5321.2045

33. Priest DG, Solano A, Lou J, Hinde E (2019) Fluorescence fluctuation spectroscopy: an invaluable microscopy tool for uncovering the biophysical rules for navigating the nuclear landscape. Biochem Soc Trans 47: 1117–1129 Doi 10.1042/bst20180604

34. Ray S, Singh N, Kumar R, Patel K, Pandey S, Datta D, Mahato J, Panigrahi R, Navalkar A, Mehra Set al (2020) α-Synuclein aggregation nucleates through liquid-liquid phase separation. Nat Chem 12: 705–716 Doi 10.1038/s41557-020-0465-9

35. Rösener NS, Gremer L, Wördehoff MM, Kupreichyk T, Etzkorn M, Neudecker P, Hoyer W (2020) Clustering of human prion protein and α-synuclein oligomers requires the prion protein N-terminus. Commun Biol 3: 365 Doi 10.1038/s42003-020-1085-z

36. Rossi M, Candelise N, Baiardi S, Capellari S, Giannini G, Orrù CD, Antelmi E, Mammana A, Hughson AG, Calandra-Buonaura Get al (2020) Ultrasensitive RT-QuIC assay with high sensitivity and specificity for Lewy body-associated synucleinopathies. Acta Neuropathol 140: 49–62 Doi 10.1007/s00401-020-02160-8

37. Ruesink H, Reimer L, Gregersen E, Moeller A, Betzer C, Jensen PH (2019) Stabilization of α-synuclein oligomers using formaldehyde. PLoS One 14: e0216764 Doi 10.1371/journal.pone.0216764

38. Ruggeri FS, Benedetti F, Knowles TPJ, Lashuel HA, Sekatskii S, Dietler G (2018) Identification and nanomechanical characterization of the fundamental single-strand protofilaments of amyloid α-synuclein fibrils. Proc Natl Acad Sci U S A 115: 7230–7235 Doi 10.1073/pnas.1721220115

39. Sano K, Atarashi R, Satoh K, Ishibashi D, Nakagaki T, Iwasaki Y, Yoshida M, Murayama S, Mishima K, Nishida N (2018) Prion-Like Seeding of Misfolded α-Synuclein in the Brains of Dementia with Lewy Body Patients in RT-QUIC. Mol Neurobiol 55: 3916–3930 Doi 10.1007/s12035-017-0624-1

40. Shahnawaz M, Mukherjee A, Pritzkow S, Mendez N, Rabadia P, Liu X, Hu B, Schmeichel A, Singer W, Wu Get al (2020) Discriminating α-synuclein strains in Parkinson’s disease and multiple system atrophy. Nature 578: 273–277 Doi 10.1038/s41586-020-1984-7

41. Singh S, DeMarco ML (2020) In Vitro Conversion Assays Diagnostic for Neurodegenerative Proteinopathies. J Appl Lab Med 5: 142–157 Doi 10.1373/jalm.2019.029801

42. Skamris T, Marasini C, Madsen KL, Foderà V, Vestergaard B (2019) Early Stage Alpha-Synuclein Amyloid Fibrils are Reservoirs of Membrane-Binding Species. Scientific reports 9: 1733–1733 Doi 10.1038/s41598-018-38271-2

43. Sorrentino ZA, Giasson BI (2020) The emerging role of α-synuclein truncation in aggregation and disease. J Biol Chem 295: 10224–10244 Doi 10.1074/jbc.REV120.011743

44. Spillantini MG, Crowther RA, Jakes R, Hasegawa M, Goedert M (1998) alpha-Synuclein in filamentous inclusions of Lewy bodies from Parkinson’s disease and dementia with lewy bodies. Proc Natl Acad Sci U S A 95: 6469–6473 Doi 10.1073/pnas.95.11.6469

45. Talens S, Leebeek FWG, Veerhuis R, Rijken DC (2019) Decoration of Fibrin with Extracellular Chaperones. Thromb Haemost 119: 1624–1631 Doi 10.1055/s-0039-1693701

46. Whiten DR, Cox D, Horrocks MH, Taylor CG, De S, Flagmeier P, Tosatto L, Kumita JR, Ecroyd H, Dobson CMet al (2018) Single-Molecule Characterization of the Interactions between Extracellular Chaperones and Toxic α-Synuclein Oligomers. Cell Rep 23: 3492–3500 Doi 10.1016/j.celrep.2018.05.074

47. Winner B, Jappelli R, Maji SK, Desplats PA, Boyer L, Aigner S, Hetzer C, Loher T, Vilar M, Campioni Set al (2011) In vivo demonstration that alpha-synuclein oligomers are toxic. Proc Natl Acad Sci U S A 108: 4194–4199 Doi 10.1073/pnas.1100976108

48. Yap TL, Velayati A, Sidransky E, Lee JC (2013) Membrane-bound α-synuclein interacts with glucocerebrosidase and inhibits enzyme activity. Mol Genet Metab 108: 56–64 Doi 10.1016/j.ymgme.2012.11.010

49. Zhou L, Kurouski D (2020) Structural Characterization of Individual α-Synuclein Oligomers Formed at Different Stages of Protein Aggregation by Atomic Force Microscopy-Infrared Spectroscopy. Anal Chem 92: 6806–6810 Doi 10.1021/acs.analchem.0c00593

50. Zucchelli S, Codrich M, Marcuzzi F, Pinto M, Vilotti S, Biagioli M, Ferrer I, Gustincich S (2010) TRAF6 promotes atypical ubiquitination of mutant DJ-1 and alpha-synuclein and is localized to Lewy bodies in sporadic Parkinson’s disease brains. Hum Mol Genet 19: 3759–3770 Doi 10.1093/hmg/ddq290

