## Supplementary Information for "Single molecule fingerprinting reveals different amplification properties of α-synuclein oligomers and preformed fibrils in seeding assay"

**Supporting figures**

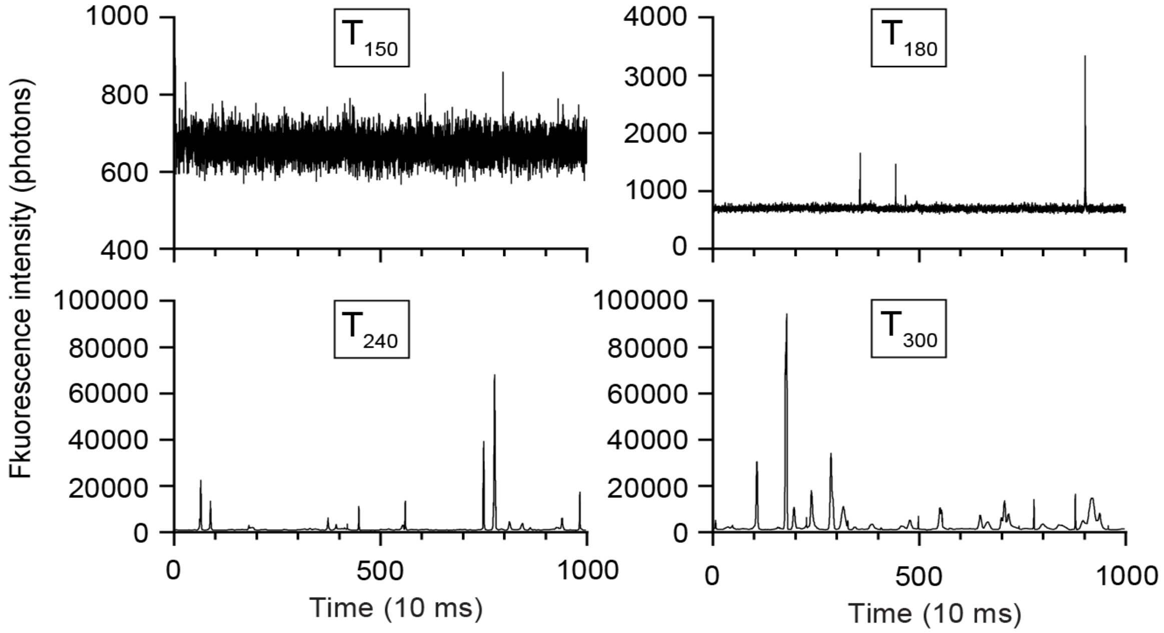


**Supporting Figure 1. Single Molecule traces of time-course aggregation experiment.**

Representative fluorescence traces during time course synthesis of oligomers and fibrils in PBS containing ThT.

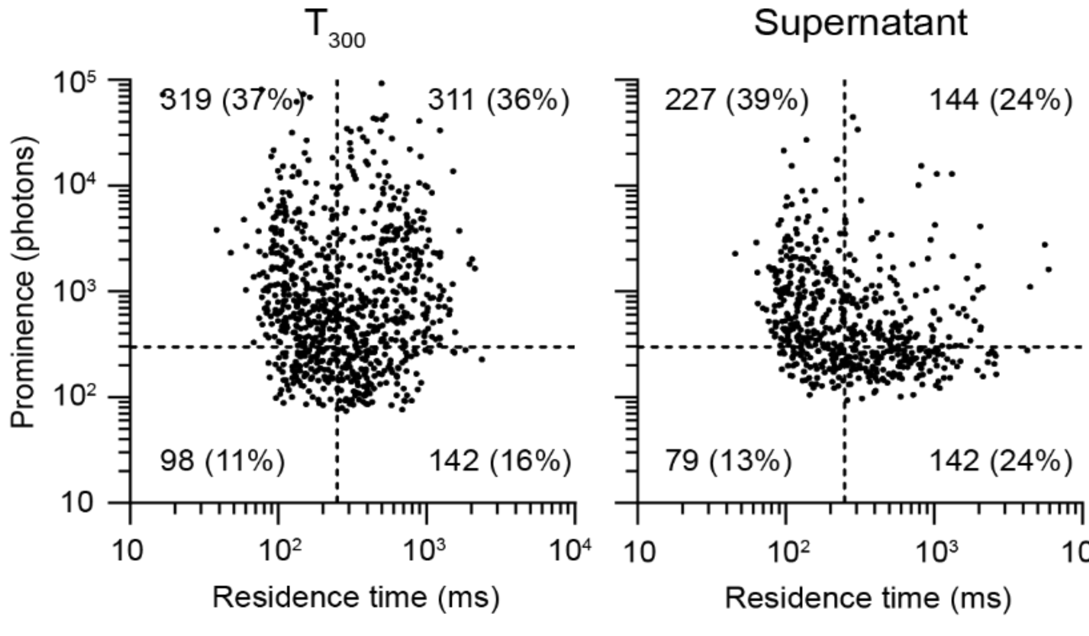


**Supporting Figure 2. Single molecule profiling of aggregation reaction and supernatant.**

The prominence and residence time of ThT events of supernatant and assembly reaction after 5 h were plotted from by pooling five different batches of supernatants measured at 20 and 200-fold dilutions. Scatter plot of the 5 h reaction is the same as **Error! Reference source not found.**B.

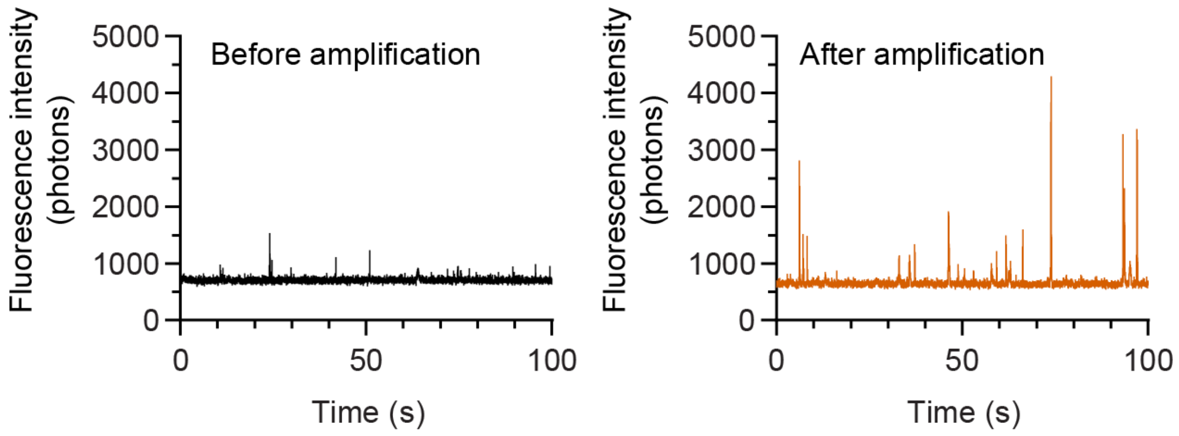


**Supporting Figure 3. Effects of amplification on oligomers.**

Representative fluorescence traces of the oligomers SEC fraction before and after amplification with human α-synuclein WT for 5 hours at 55°C.

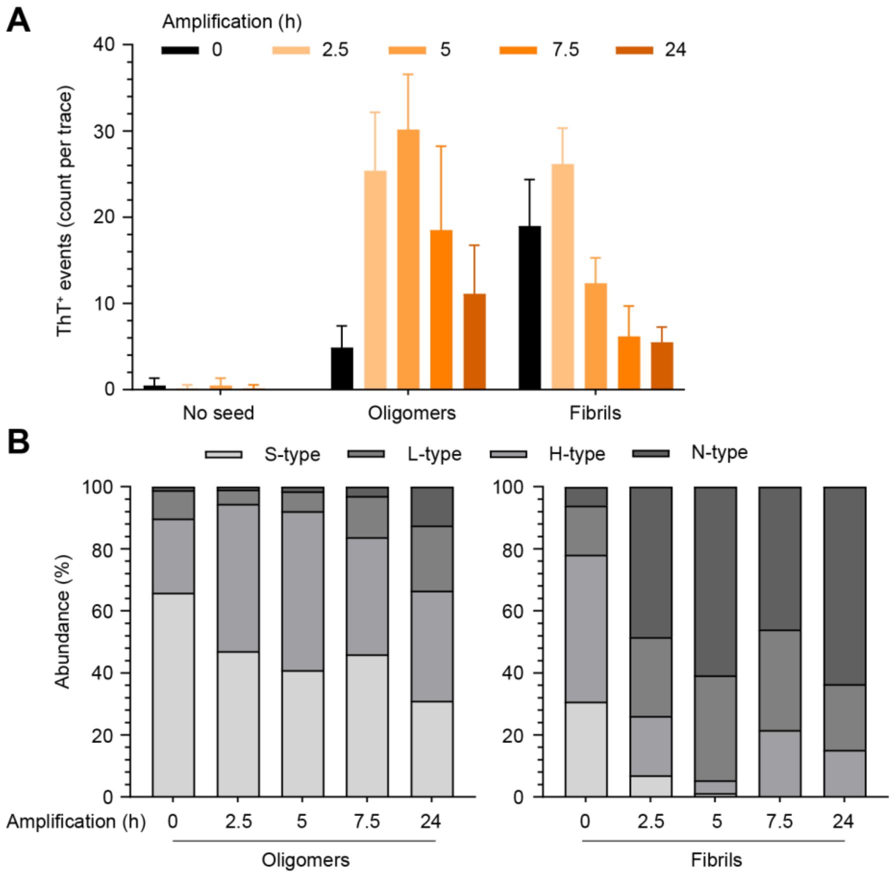


**Supporting Figure 4. Distribution of α-syn species in longitudinal study.**

**A.** Bar graphs quantifying the number of ThT+ events detected when unseeded (negative control), seeded with α-syn oligomers or PFFs. B. Bar graphs summarising the distribution of detected species from the amplification experiments in Error! Reference source not found.**A**.

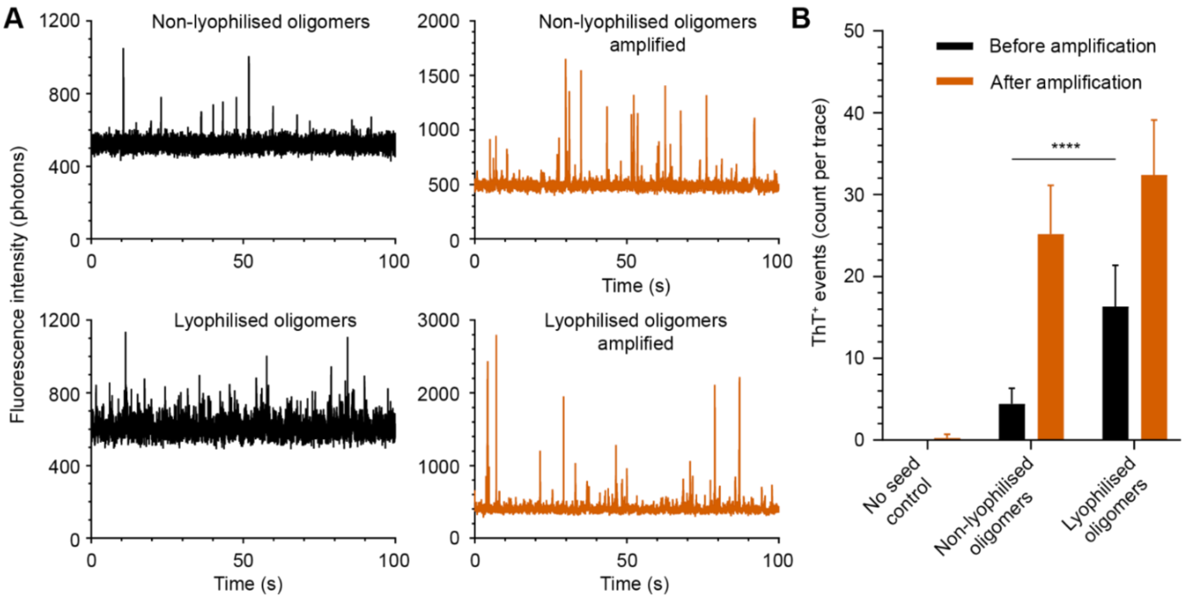


**Supporting Figure 5. Effects of lyophilisation on purified oligomers. A.** Fluorescence trace of oligomers before (black) and after amplification (red) examining the effect of lyophilisation. **B.** Bar graph of the number of ThT$^+$ events detected in non-lyophilised and lyophilised oligomers before and after amplification. Data were collected from seeding competent oligomers generated from two independent aggregation reactions. Statistics used Welch's t-test, $p \leq 0.0001$ (****).

### <u>Supporting results</u>

**Lyophilisation decreased the seeding potential of oligomers.** Two SEC eluted fractions from two independent aggregation reactions containing seeding competent oligomers in PBS were lyophilised with the LyoQuest freeze dryer (Telstar). The lyophilised products were reconstituted by adding water and mixed with α-syn WT (30 µM) and ThT (10 µM). Respective non-lyophilised oligomers were also prepared similarly, and all solutions were incubated at 55°C for 5 h for single molecule fingerprinting measurements. We observed that lyophilisation induced the formation of additional ThT$^+$ species in solution and made some oligomers less seeding competent as the change of ThT$^+$ was reduced from 5-fold to a 2-fold change (Supporting Figure 5).

### Single Molecule analysis script

```
###############################################################################
import matplotlib.pyplot as plt
import numpy as np
import matplotlib.image as mpimg
import os
from skimage import exposure
from scipy.ndimage import gaussian_filter
from skimage.morphology import reconstruction
import pandas as pd
# import cv2
from skimage import color
from os import chdir
import scipy.signal as signal
from scipy.signal import butter,filtfilt
from matplotlib.colors import LogNorm
import re
###############################################################################
c = os.getcwd()
repertoire = os.walk(c)
liste_elem = []
for root, dirs, files in repertoire :
    liste_elem.append([root, files])

###############################################################################
def butter_lowpass_filter(data, cutoff, fs, order):
    normal_cutoff = cutoff / nyq
    # Get the filter coefficients
    b, a = butter(order, normal_cutoff, btype='low', analog=False)
    y = filtfilt(b, a, data)
    return y

###############################################################################
##Step 1 : Define the filter requirements
# Filter requirements.
T = 3000          # Sample Period
fs = 1000.0       # sample rate, Hz
cutoff = 70     # desired cutoff frequency of the filter, Hz
nyq = 0.5 * fs  # Nyquist Frequency
order = 2       # sin wave
n = int(T * fs) # total number of samples
rapport = 5
ref1 = 0
taille = 120  # Calcul ecart type
mean_intensity = []
mean_width = []
mean_intengral = []
liste_nbr_peak = []
liste_name = []
liste_rapp = []
liste_mean_fin = []
#Step 2 : Filter implementation using scipy
for elem in liste_elem :
    chdir(elem[0])
    liste_fichier = elem[1]
    for fichier in liste_fichier :
        if fichier[len(fichier)-5:] != ".xlsx" and fichier[len(fichier)-4:] != ".png" and fichier[len(fichier)-4:] != ".csv" and
fichier[len(fichier)-3:] != ".py" and fichier[len(fichier)-4:] != ".txt":
            liste = pd.read_table(fichier)
            label = liste.columns[0]
            liste_y = liste[label].tolist()
            y = butter_lowpass_filter(liste_y, cutoff, fs, order)
            liste_x = np.linspace(0,len(liste_y)-1,len(liste_y))
            liste_prom = []
            liste_mean = []
```

```
for i in range (1,len(y)-taille,taille):
    liste_prom.append(np.std(liste_y[i:i + taille])*rapport)
    liste_mean.append(np.mean(liste_y[i:i + taille]))
prominence_min = min(liste_prom)
back = liste_mean[liste_prom.index(prominence_min)]
liste_mean_fin.append(back)
bruit = back/(prominence_min/rapport)**2
peaks, properties = signal.find_peaks(y, prominence =(prominence_min,None))
widths = signal.peak_widths(y, peaks, rel_height=0.5)
liste_p = []
for p in peaks :
    liste_p.append(y[p])
liste_left = widths[2]
liste_right = widths[3]
liste_intensite = []
liste_integrale = []
for i in range(len(peaks)) :
    left = int(liste_left[i])
    right = int(liste_right[i])
    liste_integrale.append(sum(liste_y[left:right])-len(liste_y[left:right])*back)
    liste_intensite.append(max(liste_y[left:right])-back)
fig = plt.figure(figsize=(50,30))
plt.plot(liste_x,liste_y, label = "prominence : " + str(prominence_min))
plt.plot(liste_x,y, label = "background : " + str(back))
plt.legend(fontsize = 60)
plt.ylabel("Intensity", fontsize = 60)
plt.xlabel("Time (10 ms)", fontsize = 60)
plt.title(fichier, fontsize = 80)
plt.tick_params(labelsize = 40)
if bruit >= ref1 :
    plt.plot(peaks, liste_p, 'o', markeredgecolor = "r", markersize = 20, label = "rapport : " + str(bruit))
    plt.legend(fontsize = 60)
    data = [[len(peaks)],
        peaks,
        liste_intensite,
        widths[0],
        liste_integrale,
        [back],
        [bruit]]
    df = pd.DataFrame(data, index = ['Peak number',
                                     'Peak position',
                                     'Peak intensity',
                                     'Peak width',
                                     'Peak intengrale',
                                     'Background',
                                     'Rapport'])
    if len(peaks)>0 :
        mean_intensity.append(sum(liste_intensite)/len(peaks))
        mean_width.append(sum(widths[0])/len(peaks))
        mean_intengral.append(sum(liste_integrale)/len(peaks))
        liste_nbr_peak.append(len(peaks))
    else :
        mean_intensity.append(None)
        mean_width.append(None)
        mean_intengral.append(None)
        liste_nbr_peak.append(len(peaks))

else :
    plt.axhline(np.mean(liste_y), linestyle= '--', color = 'r', label = 'Mean ='+ str(np.mean(liste_y)))
    plt.legend(fontsize = 60)
    data = [[np.mean(liste_y)], [prominence_min/rapport], [sum(liste_y)], [bruit]]
    df = pd.DataFrame(data, index = ['Mean',
                                     'Std',
                                     'Integral',
                                     'Rapport'])
    mean_intensity.append(np.mean(liste_y))
    mean_width.append(None)
```

```
            mean_intengral.append(sum(liste_y))
            liste_nbr_peak.append("High concentration")

        liste_rapp.append(bruit)
        liste_name.append(fichier)
        df.to_csv(fichier+".csv")
        plt.savefig(fichier+".png")
        plt.show()

chdir(liste_elem[0][0])
data = {'Name' : liste_name,
        'Peak number' : liste_nbr_peak,
        'Mean peak intensity' : mean_intensity,
        'Mean peak width' : mean_width,
        'Mean peak intengrale' : mean_intengral,
        'background' : liste_mean_fin,
        'Rapport' : liste_rapp}

df = pd.DataFrame(data)
df.to_csv("final_analyse.csv")

liste_fich = np.linspace(0,len(liste_rapp)-1,len(liste_rapp))

fig = plt.figure(figsize=(50,30))
plt.plot(liste_fich,liste_rapp, 'o', markeredgecolor = "r", markersize = 30)
plt.axhline(ref1, linestyle= '--', color = 'b', label = 'rapport ='+ str(ref1))
plt.legend(fontsize = 60)
plt.ylabel("Rapport", fontsize = 60)
plt.xlabel("Files", fontsize = 60)
plt.title("Rapport", fontsize = 80)
plt.tick_params(labelsize = 40)
plt.savefig("Rapport.png")
plt.show()

fig = plt.figure(figsize=(50,30))
plt.plot(liste_fich,liste_mean_fin, 'o', markeredgecolor = "r", markersize = 30)
plt.legend(fontsize = 60)
plt.ylabel("Mean", fontsize = 60)
plt.xlabel("Files", fontsize = 60)
plt.title("Mean", fontsize = 80)
plt.yscale("log")
plt.tick_params(labelsize = 40)
plt.savefig("Mean.png")
plt.show()
```
